# Taxonomically Restricted Genes are Associated with Responses to Biotic and Abiotic Stresses in Sugarcane (Saccharum spp.)

**DOI:** 10.1101/2022.04.29.489768

**Authors:** Claudio Benicio Cardoso-Silva, Alexandre Hild Aono, Melina Cristina Mancini, Danilo Augusto Sforca, Carla Cristina da Silva, Luciana Rossini Pinto, Keith L. Adams, Anete Pereira de Souza

## Abstract

Orphan genes (OGs) are protein-coding genes that are restricted to particular clades or species and lack homology with genes from other organisms, making their biological function difficult to predict. OGs can rapidly originate and become functional; consequently, they may support rapid adaptation to environmental changes. Extensive spread of mobile elements, and whole genome duplication, occurred in the *Saccharum* group, which may have contributed to the origin and diversification of OGs in the sugarcane genome. Here, we identified and characterized OGs in sugarcane, examined their expression profiles across tissues and genotypes, and investigated their regulation under varying conditions. We identified 319 OGs in the *Saccharum spontaneum* genome without detected homology to protein-coding genes in green plants, except those belonging to Saccharinae. Transcriptomic analysis showed 288 sugarcane OGs with detectable expression levels in at least one tissue or genotype. We observed similar expression patterns of OGs in sugarcane genotypes originating from the closest geographical locations. We also observed tissue-specific expression of some OGs, possibly indicating a complex regulatory process for maintaining diverse functional activity of these genes across sugarcane tissues and genotypes. Sixty-six OGs were differentially expressed under stress conditions, especially cold and osmotic stresses. Gene co-expression network and functional enrichment analyses suggested that sugarcane OGs may be involved in several biological mechanisms, including stimulus response and defence mechanisms. These findings provide a valuable genomic resource for sugarcane researchers, especially those interested in selecting stress-responsive genes.

## 1 INTRODUCTION

Recent advances in sugarcane genomics have created opportunities to systematically understand the evolutionary history and diversification of the *Saccharum* group. However, the complexity of the sugarcane genome, mainly due to its size, ploidy level, and the large number of mobile elements (Thirugnanasambandam et al., 2018), has hindered advances in the genomic field of this important crop species. Despite the economic importance of sugarcane due to its use as a source of sugar, biofuel, and fibre, its reference genomes, including a chromosome-level *Saccharum spontaneum* genome (Zhang et al., 2018), a monoploid genome from the R570 variety (Garsmeur et al., 2018), and an SP80-3280 hybrid genome (Souza et al., 2019), have been only recently reported.

It has been suggested that two events of whole genome duplication (WGD) occurred during the evolution of the Saccharum group (Ming et al., 1998; Paterson et al., 2012). These recent events of polyploidization occurred within Saccharinae group provide an opportunity to investigate the fate of duplicated genes. WGD is a major mechanism responsible for species diversification and adaptation (Soltis et al., 2009; Renny-Byfield and Wendel, 2014). Genome duplication initially results in gene duplication and gene redundancy. After duplication, some gene copies preserve their original function, while most of them are eliminated through negative selection (Tautz and Domazet-Lošo, 2011). However, some copies under positive selection, after sequence diversification, may acquire a new biological function (Peer et al., 2009). Divergence of pre-existing genes is one of the mechanisms underlying the emergence of new genes (Tautz and Domazet-Lošo, 2011).

Taxonomically restricted or orphan genes (OGs), which have no homology to genes in other taxa, may contribute to evolutionary novelties and might be responsible for lineage-specific trait origins (Wilson et al., 2005; Khalturin et al., 2009; Tautz and Domazet-Lošo, 2011). Previous comparative genomic studies have estimated that OGs constitute at least 1% of the total genes in a genome, depending on the alignment rate and taxonomic level considered (Khalturin et al., 2009; Arendsee et al., 2014; Prabh and Rödelsperger, 2016). Comparative genomics applied to investigate OGs has demonstrated that these genes have shorter lengths and intron sizes and that they have higher levels of transposable elements than genes broadly shared across green plants (Tautz and Domazet-Lošo, 2011).

Previous studies described OGs as being regulated in response to environmental changes, especially those associated with biotic and abiotic stresses (Beike et al., 2015; Giarola et al., 2015; Khraiwesh et al., 2015; Schlötterer, 2015). For example, some taxonomically restricted genes were found to be differentially expressed (DE) in tomatoes infected with viruses (Kaur et al., 2017). In Arabidopsis, the well-studied OG *QQS* (*Qua-Quine Starch*) is involved in the regulation of nitrogen allocation, and when *QQS* is transferred and expressed in other species, it affects important traits such as protein content, consequently increasing crop yields (O’Conner et al., 2018). Previous works also reported that OGs play a vital role in soluble sugar metabolism in *Brassica rapa* based on an in-depth analysis of orphan genes by establishing a gene editing system and phenotypic validation (Jiang et al., 2020).

Despite the biological relevance of these taxonomically restricted genes, there is no previous report describing their occurrence and expression patterns in the Saccharum complex. In this study, we identified and characterized sugarcane orphan genes and their expression patterns across tissues and genotypes. Additionally, we tested under which conditions these genes are positively or negatively regulated.

## 2 MATERIALS AND METHODS

### 2.1 Orphan Gene Identification

A phylostratigraphic approach based on a sequence homology search was used to identify orphan genes in the sugarcane genome (Domazet-Lošo et al., 2007; McLysaght and Hurst, 2016). These analyses rely on sequence alignments to detect genes that lack homology in a focal species in comparison to a target clade. For this analysis, we used the gene model from *S. spontaneum* (Zhang et al., 2018) as a reference. First, the protein and coding DNA sequence (CDS) files containing the set of sugarcane genes were filtered using the CD-HIT package v.4.8.1 (Fu et al., 2012); a similarity threshold of 90% was applied for both the CD-HIT and CD-HIT-EST algorithms, which were employed for the protein and CDS files, respectively. This step was performed to remove redundancies in the dataset once all the homologous genes and duplications were included in the annotated sugarcane genome. Subsequently, a series of alignments using both the sugarcane proteome and CDSs were performed using the BLAST algorithm with e-value ≤ 1*e*^−6^ to identify sugarcane orphan genes. In this step, CDSs and protein sequences from Viridiplantae species (*Arabidopsis thaliana* TAIR10, *Brachypodium distachyon* v3.1, *Citrus sinensis* v3.1, *Eucalyptus grandis* v2.0, *Miscanthus sinensis* 7.1, *Oryza sativa* 7.0, *Phaseolus vulgaris* v2.1, *Panicum virgatum* v4.1, *Setaria italica* v2.2, *Solanum lycopersicum* ITAG3.2, *Sorghum bicolor* v3.1.1, *Oropetium thomaeum* v1.0, and *Zea mays* 284 v6) were downloaded from Phytozome v.13 (Goodstein et al., 2012) and converted into a database to perform the alignments. All the sugarcane genes with detected homologies to other genes from these target species were filtered out. The remaining subset of sugarcane genes was aligned to non-redundant protein (NR) and nucleotide (NT) NCBI databases, and genes with identified homologs not included in previous filtering steps were discarded. We used the same BLAST parameters as in the first step of the orphan genes filtering.

### 2.2 Orphan Gene Duplication and Manual Curation of Homeologs

Because OGs lack homology with genes from other plant species, we checked whether some OG duplicates were missed across sugarcane chromosomes. Then, OG CDSs were mapped back onto the sugarcane chromosomes using sim4 software (Florea et al., 1998). The coordinate position of each putative OG exon was used as a starting point for manual curation, and Artemis v.18.0 (Carver et al., 2012) was used to check intron/exon boundaries as well as the presence of start and stop codons. The FASTA files containing sugarcane chromosome information and gff3 files were used for the manual curation of the OGs. Transposable elements in the orphan gene sequences were predicted using a reference collection of repeats from the Repbase database, which includes grass family elements (Bao et al., 2015).

### 2.3 RNA-Seq Experimental Data: Retrieving and Preprocessing

An extensive search for sugarcane papers reporting RNA-Seq data was performed, followed by a search of the SRA NCBI repository. The selected RNA-Seq samples were retrieved using the ‘fastq-dump’ program from the SRA toolkit (version 8.22), and SRA files were converted to fastq-format files. The raw reads were subjected to quality control using Trimmomatic v0.36 (Bolger et al., 2014) to remove adapter and low-quality sequences. Three reference transcriptomes, including two full-length transcriptomes, were also selected to confirm which selected OGs were being transcribed. In the first set of IsoSeq data, RNA samples were extracted from the top and bottom internodes of 22 genotypes (Hoang et al., 2017), and in the second, RNA samples were obtained from leaves of the commercial sugarcane variety from Thailand (Piriyapongsa et al., 2018). The third transcriptome, which was de novo assembled from short reads, was extracted from the leaves of six sugarcane hybrids (Cardoso-Silva et al., 2014). A local alignment using the BLASTn program was performed with an e-value cut-off ≤1*e*^−6^ to identify the homology of putative orphan genes to sugarcane transcripts.

### 2.4 Orphan Gene Expression Profile and Differential Expression

RNA-Seq libraries were selected for two purposes: (*i*) to unveil expression patterns of OGs across sugarcane tissues and genotypes and (*ii*) to identify DE OGs, especially under stress conditions. The expression level of each gene was estimated by mapping the transcriptomes against the whole gene set of sugarcane using Salmon (Patro et al., 2017). The level of expression of a given gene was calculated by the log transform method implemented in Salmon (transcripts per million (TPM)), which represents the relative abundance of a transcript among a population of transcripts. Heatmaps representing the expression levels of OGs in both differential gene expression and expression across tissues/genotypes experiments were built using the superheat R package (Barter and Yu, 2018).

To investigate whether OGs are DE, we selected RNA-Seq experiments including biotic and abiotic stresses, developmental stages, and sucrose accumulation. Cleaned reads from each library originating from experiments with biological replicates were mapped to the complete set of sugarcane genes (CDS FASTA format) using Salmon v.0.12.0 to quantify the transcript abundance (Patro et al., 2017). DESeq2 package v.3.9 (Love et al., 2014) was used to predict differentially expressed genes (DEGs) in each experiment using the raw read counts as input data. In cases where the same sample was sequenced in multiple runs, the technical replicates were collapsed before starting the DEG analysis. To minimize quantification biases, genes with less than 10 reads mapped per sample were filtered out before the gene expression analysis. The DEGs were estimated assuming a negative binomial distribution for each gene, applying a function that estimates the size factor and reducing bias caused by library size (normalization by median of rations; gene count divided by sample size factor). A *p-value <* 0.05 and absolute log_2_ Fold Change ≥ 2 were used as thresholds for determining whether genes were apparently DE. For each predicted OG, a hypothesis was tested to determine whether there was enough evidence against the null hypothesis that there were no differences between the control and treatment groups, thus supporting the assumption that any difference in gene expression occurred merely by chance.

### 2.5 Gene Co-Expression Network

Co-expression networks were built to detect modules enriched with OGs. The expression matrix of the 218 samples across sugarcane tissues and genotypes was used to build a co-expression network with the WGCNA R package (Langfelder and Horvath, 2008). A weighted adjacency matrix was constructed using pairwise Pearson’s correlation coefficient measures and an estimated power threshold to fit a scale-free independence (*R*^2^ *>* 0.8 and largest mean connectivity). Subsequently, the calculated matrix was converted to a topological overlap matrix (TOM), which evaluates gene pair correlations and the degree of agreement with other genes in the matrix (Yip and Horvath, 2007). After that, we estimated network functional modules by employing average linkage hierarchical clustering in accordance with the TOM-based dissimilarity measure. We used a soft threshold power of 7 (*R*^2^ = 0.81 and mean connectivity 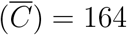) for calculating the TOM matrix. Clusters were defined from a hierarchical dendrogram using adaptive branch pruning implemented in the dynamicTreeCut R package (Langfelder et al., 2008).

## 3 RESULTS

### 3.1 Detection of Sugarcane Orphan Genes

After removing redundancies in the sugarcane gene model, which contained 83, 826 genes from *S. spontaneum* (Zhang et al., 2018), we obtained a total of 51, 675 non-redundant protein coding genes. These sugarcane genes were aligned to the proteomes of thirteen grasses, represented by 435, 957 proteins. This alignment returned 1, 536 sugarcane genes with no homology with any other grass protein. Next, these subsets of ‘no-hit’ genes were aligned to the NR protein database, and 442 genes were found with no homology. Finally, these remaining sets were aligned to the NT database. As a result, a total of 335 genes were identified as sugarcane orphan genes due to their lack of homology to other genes.

Homology searches may fail to detect homologous in other species, resulting in spurious orphan genes prediction. To minimize this effect, we did not rely only on homology searches, but we also mapped OGs CDS sequences onto each chromosome of seven grass genomes (*S. spontaneum, M. sinensis, S. bicolor, P. hallili, Z. mays, S. italica, O. sativa*) and *A. thaliana*. Although OGs were not annotated as complete genes, except in the Saccharum group, we found vestiges of exons of these genes in all grass chromosomes. However, we did not detect OGs vestiges in *A. thaliana* chromosomes (*Supplementary Table 2*).

To better understand the distribution of these putative OGs across grass genomes, we assessed chromosome regions in which dispersed fragments of the OGs aligned with at least 10% of their length. Intriguingly, the closer the phylogenetic relationship with *Saccharum*, the higher the number of OG fragments observed in grass genomes (*Figure 1* and *Supplementary Table 2*). The number of OG fragments ranged from 114 in the *O. sativa* genome to 91, 379 in the *M. sinensis* genome. If we consider the *S. spontaneum* genome, the number of OG fragments is even larger. Curiously, there are 25 times more OG fragments on *S. spontaneum* and *M. sinensis* chromosomes than in the sorghum genome, which is the closest relative of these Saccharinae species.

**Figure 1.**
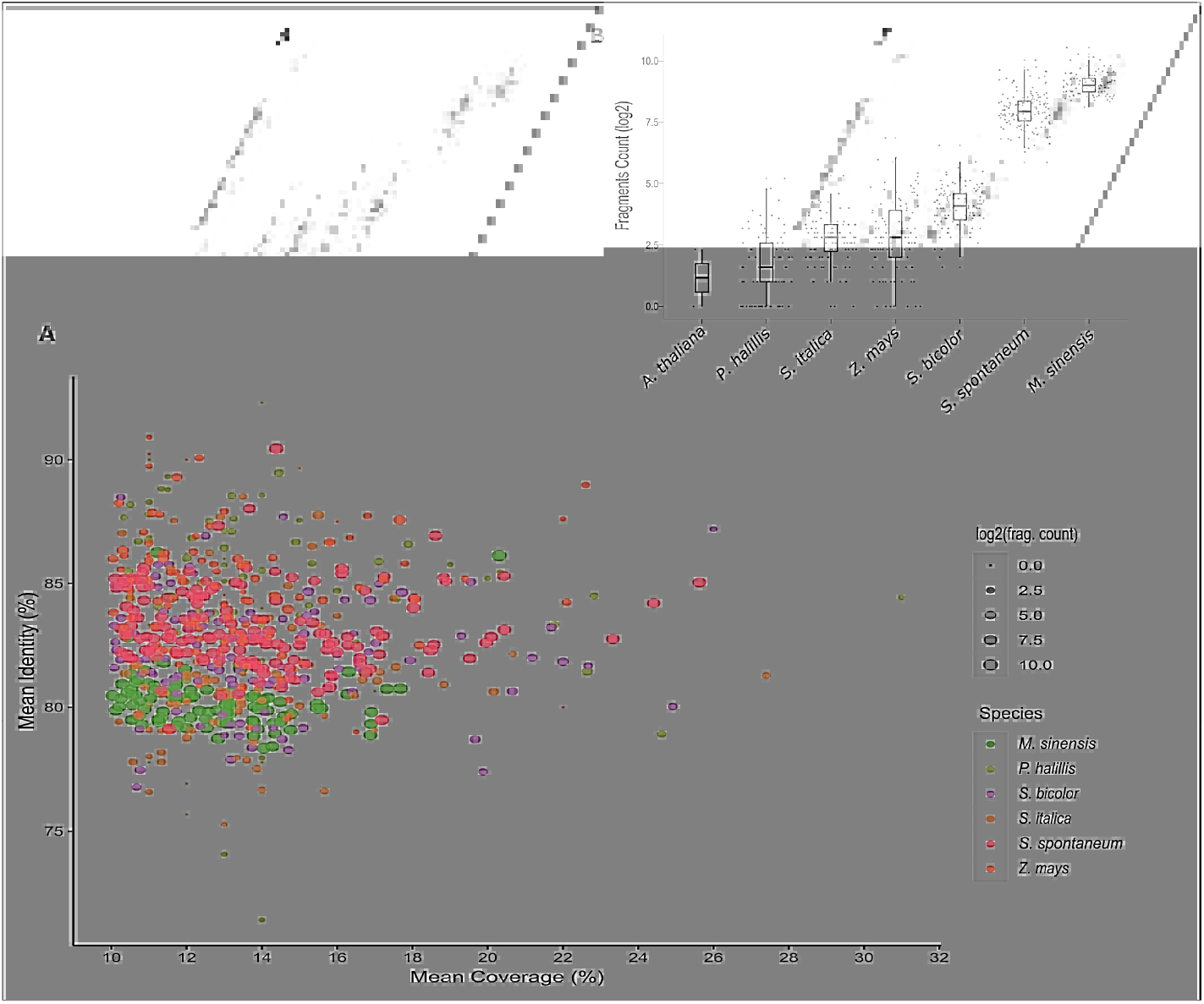
Evidence of orphan gene (OG) vestiges within grass genomes. (A) Scatterplot showing the average percentage identity and coverage of each OG in six grass species. Each dot represents an individual OG. The dot sizes represent the number of fragments of each sugarcane OG in other grass species. The number of fragments of each OG in the focal species is also shown in the bar plot (B).

The large number of gene fragments found in the Saccharinae genomes may suggest that some of these genes originated from transposable element (TE) duplication. We aligned all the predicted OGs onto TE sequences to test the hypothesis that some of the OGs are transposable elements and to verify whether some of the OGs were derived from TE insertion (*Supplementary Table 3*). This search was motivated by previous observations suggesting that 51% of the orphan genes in rice are derived from TEs (Jin et al., 2021). To investigate this hypothesis more deeply and shed light on the origin of these putative lineage-specific genes, we performed an alignment of the initially selected set of genes (335 OGs) against the [Repbase](http://www.girinst.org/repbase) TE library. A total of 153 putative OGs aligned on TEs with significant hits (E-value ≤ 1*e*^−10^). For a subset of these genes (16 OGs), at least 70% of the sequences aligned on TEs. We assumed that these genes are putative TEs, and we did not consider them to be sugarcane OGs. We selected a final set of 319 orphan genes to test whether OGs are DE under stress conditions and to quantify their expression levels across sugarcane tissues and genotypes. These selected OGs had an average of 3.54 exons (*σ* = 2.37; ranging from 1 to 21), and 57.9% had three exons or fewer. A total of eight OGs had between 10 and 21 exons (*Supplementary Table 4*).

### 3.2 Evidence of Orphan Gene Expression across Sugarcane Tissues and Genotypes

We searched for evidence that the OGs were being transcribed across several sugarcane tissues and genotypes by aligning them against reference transcriptomes and RNA-Seq libraries. We selected two representative sugarcane IsoSeq datasets (Hoang et al., 2017; Piriyapongsa et al., 2018), a collection of transcripts from six sugarcane varieties (Cardoso-Silva et al., 2014), and 218 RNA-Seq samples from sugarcane hybrids, *S. officinarum*, and *S. spontaneum* (*Supplementary Table 1*). Evidence of transcription was detected in 89.34% of the OGs (*TPM* ≥ 1), which were expressed in at least one transcriptome experiment or tissue (*Supplementary Tables 4 and 5*). We observed that almost one-third of the OGs had an expression level considered low (1 ≥ *TPM* ≤ 10), while only 6% of them had a value greater than 100 TPM. In fact, when we compared the expression level of OGs and non-OGs in four sugarcane tissues (*Supplementary Figure 1*), we found that OGs have proportionally lower expression levels. However, these differences were almost imperceptible in the meristem tissue (bud).

The expression profile of OGs across tissues and genotypes revealed a clear pattern of clustering of the samples. Overall, OGs showed a similar expression pattern among genotypes when we compared the same tissue (*Figure 2* and *Supplementary Table 5*). Although the expression matrix combined tissues and genotypes, we also observed a clustering profile by genotype to the same degree. For example, samples originating from *S. spontaneum* (Krakatau, IN84_58, and SES205A) were clustered together and separated from those originating from *Saccharum officinarum* (BadilaDeJava, CriollaRayada, WhiteTransparent) and *Saccharum* hybrids. Notably, samples originating from *Saccharum* hybrids and *S. officinarum* had more similar expression patterns than those originating from *S. spontaneum*.

**Figure 2.**
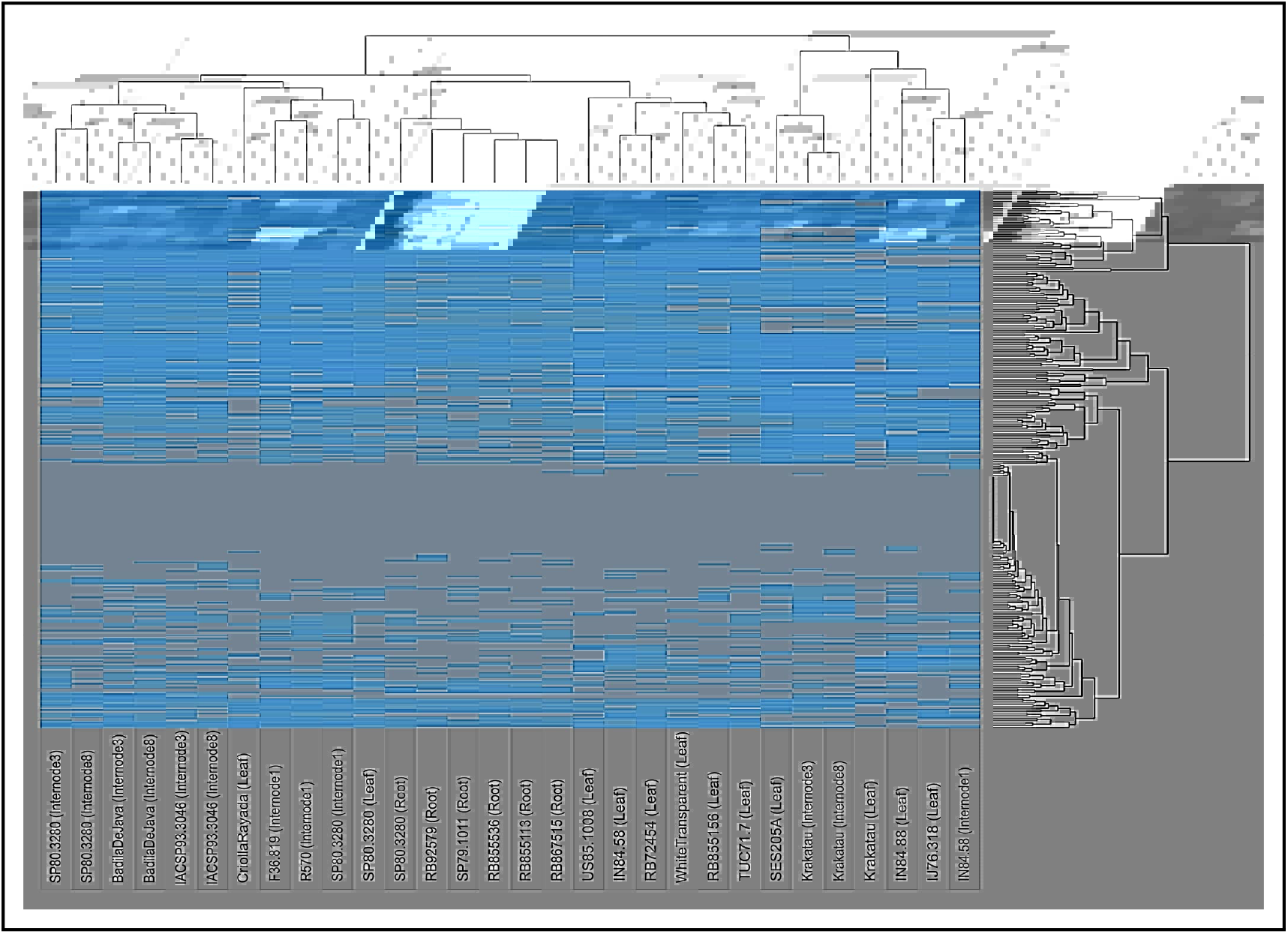
Expression profiles of orphan genes (OGs) in several sugarcane tissues and genotypes. The expression of each gene (TPM) was estimated using a pseudoalignment method implemented in Salmon software (Patro et al., 2017)

In particular, the expression pattern of OGs in the internodes of *S. spontaneum* was noticeably different when compared to that in *S. officinarum* and hybrids. Furthermore, some OGs present a contrasting expression pattern when comparing samples from *S. officinarum* and *S. spontaneum*. Specifically, a subset of 42 OGs had an average expression level that was three times higher in *S. officinarum* than in *S. spontaneum*. In contrast, the expression of 35 OGs was higher in *S. spontaneum*. Interestingly, we also observed that genotypes originating from the same geographical location tended to be clustered together because they had more similar expression profiles. In particular, this pattern was observed in the expression levels of OGs in internode 1 of French hybrids (F36.819 and R570) and roots of Brazilian hybrids (RB855536, RB855113, and RB867515).

Interestingly, a few OGs seem to have tissue-specific regulation, as observed for some genes only expressed in roots (Sspon.06G0001310 and Sspon.06G0024430), internode 1 (Sspon.08G0030340), internodes 3 and 8 (Sspon.05G0032460 and Sspon.08G0007980), and all internodes (Sspon.06G.0027080).

### 3.3 Most Orphan Genes are Differentially Expressed Under Stress Conditions

We selected eight RNA-Seq experiments representing a variety of conditions to test whether OGs change their expression level (*Table 1*). After filtering the raw data, we selected more than 13 billion high-quality reads to perform gene expression analysis.

**Table 1.** RNA-Seq experiments selected to perform sugarcane gene expression analysis.

| Accession | Condition | Cultivar/Species | Tissue | Replicates | References |
| --- | --- | --- | --- | --- | --- |
| PRJNA474042 | Yellow Canopy Syndrome | Hybrid | leaf | 5 | (Marquardt et al., 2019) |
| PRJNA479814 | developmental stages | Q208 and KQ228 | root/leaf/internode | 3 | (Thirugnanasambandam et al., 2019) |
| PRJNA483518* | Cold Stress | Guitang08-1180 and ROC22 | leaf | 4 | (TANG et al., 2018) |
| PRJNA533093 | Low Nitrogen | Badila | leaf and root | 3 | (Yang et al., 2019) |
| PRJNA291816 | Smut Disease | RB925345 | bud | 3 | (Bedre et al., 2019) |
| PRJNA415122 | *S. scitamineum*-infected | CP74_2005 | bud | 3 | (McNeil et al., 2018) |
| PRJNA371469 | Osmotic Stress | *S. officinarum* | root and leaf | 2 | (Pereira-Santana et al., 2017) |
| PRJNA681593 | Sucrose Accumulation | Hybrids | top/bottom internodes | 3 | (Aono et al., 2021) |

\*Represented by multiple accessions (see more details in *Supplementary Table 01*).

We observed at least one OG DE in five out of eight RNA-Seq experiments (*Supplementary Table 6*). We did not detect OGs differentially expressed in the RNA-Seq experiments related to developmental stage, sucrose accumulation, and plants infected with *Sporisorium scitamineum*. Generally, more genes were DE in the experiments related to abiotic stress than in those related to biotic stress. For example, we estimated that 6, 440 genes were DE under cold stress (Guitang08-1180 and ROC22 hybrids), while only 1, 548 genes were DE after infection with yellow canopy syndrome, and 2, 612 genes were DE in plants infected with smut disease. Overall, we identified 66 OGs that were DE under at least one type of stress (log_2_ *FoldChange ≥* 2; *padj <* 0.05). Most of the genes were regulated by abiotic stresses, while only four OGs were regulated by biotic stresses. We detected only two OGs (Sspon.02G0052680-1C and Sspon.06G0009900-2C) that were DE in both osmotic stress and cold stress experiments, and one gene (Sspon.04G0024550-1B) was shared between the cold stress and low-nitrogen experiments.

In the cold stress experiment, we identified 8, 921 DE genes in the hybrid Guitang08-1180 (5, 644 upregulated and 3, 277 downregulated) and 8, 913 DE in the hybrid ROC22 (5, 596 upregulated and 3, 317 downregulated), and 72% of these genes were DE in both genotypes. A total of 50 OGs were determined to be DE in this experiment, with 33 OGs in Guitang08-1180 (12 upregulated and 21 downregulated) and 36 OGs in the ROC22 hybrid (12 upregulated and 24 downregulated), while 18 OGs were DE in both genotypes (*Figure 3*, and *Supplementary Table 6*). An OG that was upregulated in both genotypes (Sspon.06G00100300) has three conserved copies annotated in the *S. spontaneum* genome (*Supplementary Figure 2*).

**Figure 3.**
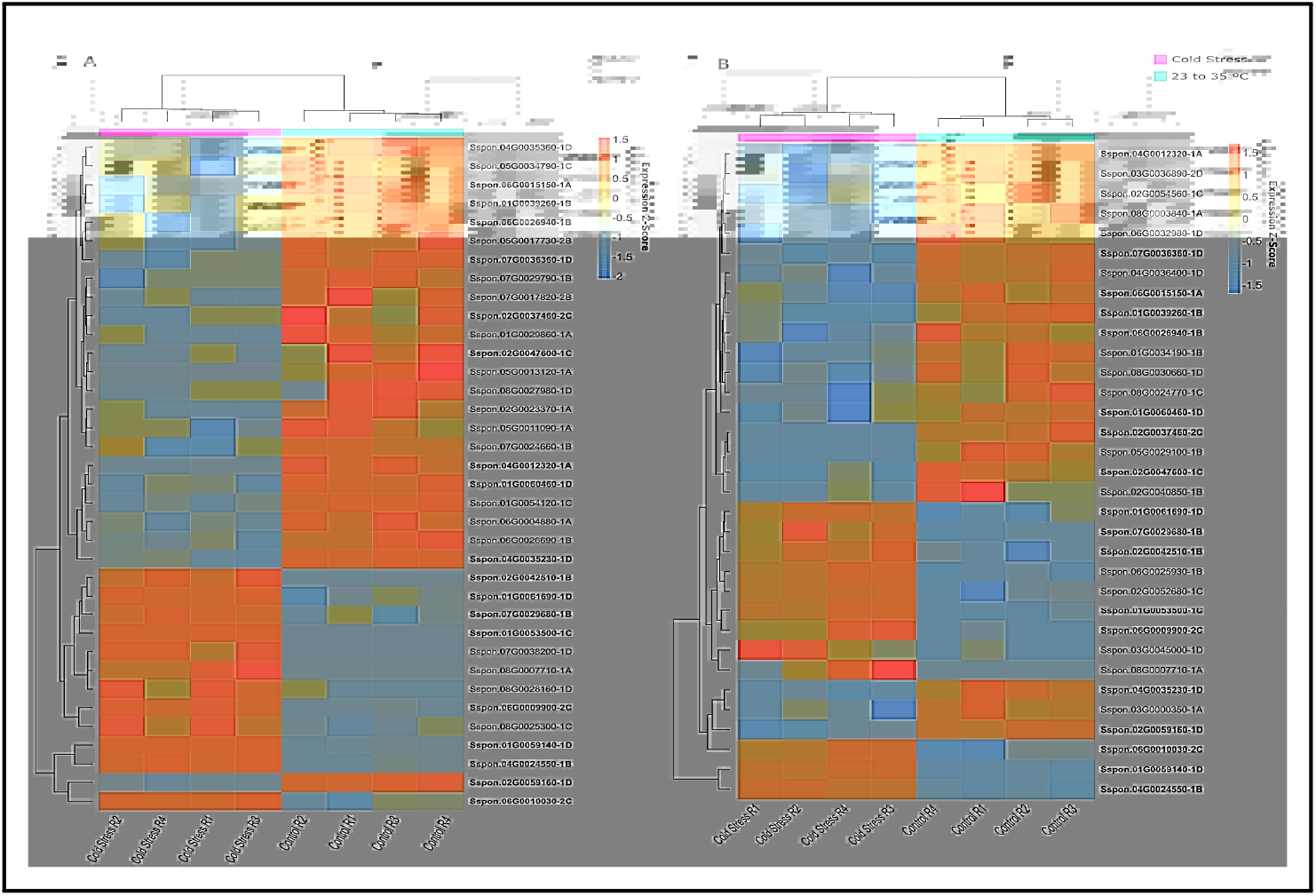
Orphan genes (OGs) differentially expressed (DE) under cold stress. Hierarchical clustering of the genes expressed at normal and cold temperatures with four replicates for each treatment (row). GE analysis was carried out in leaf tissues at an ambient temperature (ranging from 23*°C* to 35*°C*) and a cold temperature (4*°C* in a well-controlled climate chamber) in two sugarcane genotypes: ROC22 (A) and Guitang08-1180 (B).

In the osmotic stress experiment conducted on leaves and root samples from *S. officinarum*, a total of 4, 207 genes were DE in the leaves (1, 815 upregulated and 2, 392 downregulated), while 4, 222 genes were DE in the root samples (1, 707 upregulated and 2, 515 downregulated). Of these genes, 13 OGs were identified as DE, nine in the leaf samples and six in the root samples. Two OGs, Sspon.01G0060030-1D and Sspon.05G0013120-1A, were detected as DE in both the root and leaf tissues (*Figure 4, Supplementary Table 6*).

**Figure 4.**
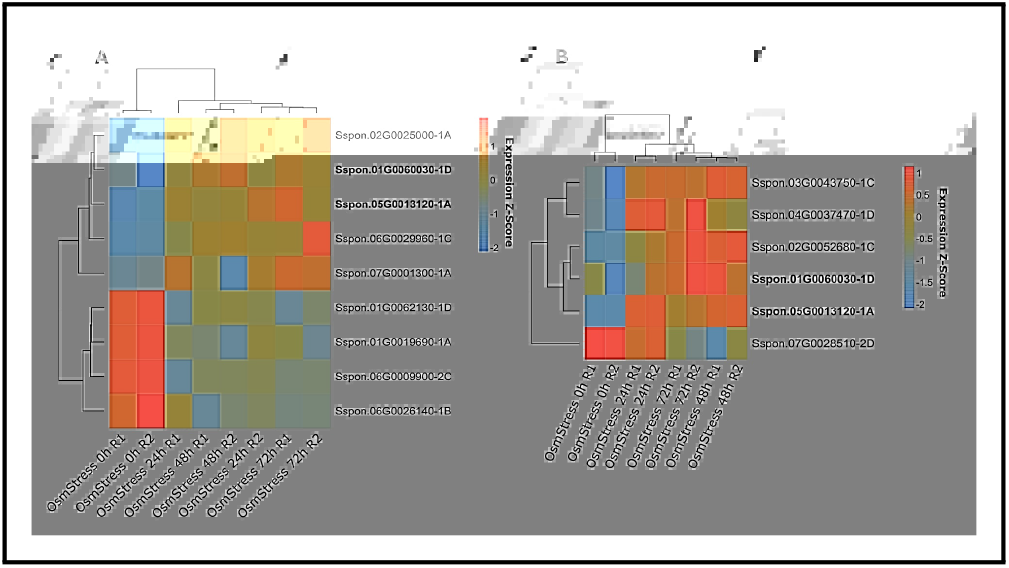
Heatmap of orphan genes (OGs) differentially expressed (DE) under osmotic stress. Plants were subjected to osmotic stress for 24, 48, and 72 h, and other plants were maintained without osmotic stress (0 h). RNA samples were extracted from the leaves (A) and roots (B) of *S. officinarum*.

**Figure 5.**
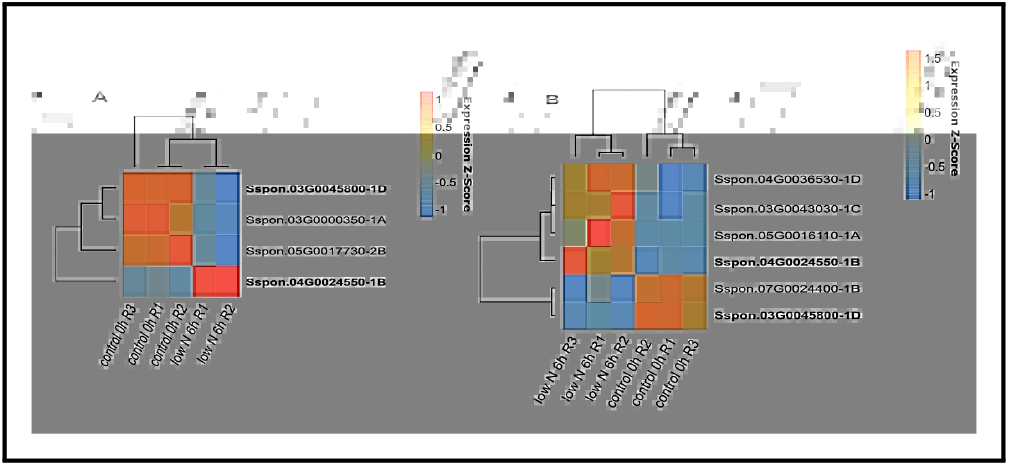
Genes differentially expressed (DE) under low-nitrogen conditions. RNA samples were extracted from leaves of the ROC22 (A) and Badila (B) genotypes.

In the sugarcane plants exposed to low-nitrogen conditions, most of the DE genes were detected in the leaf samples (4, 524 genes in the Badila variety and 2, 345 genes in the ROC22 hybrid), while in the roots, 894 and 726 genes were estimated to be DE in Badila and ROC22, respectively. In the root samples, the number of genes upregulated was three times greater than the number of genes downregulated. This pattern was observed in both genotypes. We did not detect DE OGs in the roots of both genotypes; however, in the leaves, we detected six DE genes in Badila (four upregulated and two downregulated) and four DE genes in the ROC22 hybrid (one upregulated and three downregulated) (*Figure reffig:5, Supplementary Table 6*).

### 3.4 Functional Predictions of Orphan Genes

A guilt-by-association principle was used to obtain some clues about OG functionality once it was determined that these genes had no homology to any known protein in other organisms. The co-expression network was built considering the full set of sugarcane genes, including 288 orphan genes with estimated expression values. As a result, we identified a total of 153 modules, of which 78 had at least one orphan gene included. An enrichment analysis of these modules containing OGs suggests that these genes are associated with several biological processes (*Supplementary Table 7*). Overall, the most frequent gene ontology terms observed in modules containing OGs included those associated with responses to several stimuli and defence mechanisms (*Supplementary Figure 3*). The modules that contained more orphan genes (64 OGs) was functionally enriched in terms of metabolic and biosynthesis processes as well as transport and stimulus responses. A search for protein domains within OG sequences revealed that most OGs do not have similarity with any functional domains deposited in the Pfam database. Based on these searches, we detected only partial local alignment for 30 OGs (BLAST searches produced no significant alignments to domains; E-value ≥ 1*e*^−3^), of which nine were predicted to be domains of unknown function. Interestingly, a subset OGs had partial alignment with domains associated with defence and stress tolerance (*Supplementary Table 7*).

## 4 DISCUSSION

The definition and determination of OGs is context dependent, but generally, a gene that lacks detectable sequence homology in other taxa is typically an OG (Rödelsperger et al., 2019). The fact that these OGs do not have homology with other genes in the target species raises the question of whether they could have evolved from noncoding regions, as previously reported (Oss and Carvunis, 2019; Vakirlis et al., 2020). Surprisingly, after mapping the CDSs of the OGs onto grass chromosomes, we detected fragments of them scattered throughout these genomes. The number of fragments was inversely proportional to the phylogenetic distance, where the highest number of OG fragments was observed in *M. sinensis* chromosomes, the species closest to sugarcane. A similar case study was reported in nematodes, where the authors found traces of homology with OGs in closely related genomes; the authors described in detail the de novo origin of OGs from non-coding regions (Prabh et al., 2018). This finding may indicate that some OGs rapidly emerged de novo into the Saccharinae group given that they do not have any vestiges in other grass genomes or that the fragments were amplified within Saccharinae through retrotransposition (Cerbin and Jiang, 2018). We also observed that many OG fragments were retained, independent of the phylogenetic relationship. For example, the OGs Sspon.03G0016730-2B and Sspon.04G0031650-1C have more than 85% of their CDS features covered in the *S. italica* chromosome, while their coverage in the *Z. mays* chromosome is less than 20%.

Surprisingly, we identified some OGs with lengths and exon numbers greater than those previously described for taxonomically restricted genes; OGs have small lengths, and most have only one exon (Campbell et al., 2007; Guo et al., 2007; Xu et al., 2015; Jin et al., 2021). Although most of the OGs we have identified contained three or fewer exons, some had more than 10 exons.

Sugarcane genome complexity and the lack of reference genome sequences for the hybrids and *S. officinarum* impose a layer of complexity when estimating expression level. Additionally, previous work reported that the expression levels of OGs were lower than those of non-OGs (Li et al., 2019). Expression estimation tends to be more difficult to implement and imprecise for genes with low levels of expression because a low read count leads to high Poisson noise, and any biological effect may become biased (Munsky et al., 2012). The selection of RNA-Seq experiments with at least three replicates helped to improve accuracy when predicting which genes are differentially expressed. Nevertheless, we selected a set of genes that evolved recently within the Saccharinae group, which had a short evolutionary period over which duplication and structural variation could have occurred, consequently increasing the likelihood of successful alignment.

The expression profiles of OGs across sugarcane hybrids may indicate that these genes are regulated at the same level among genotypes from the same areas. Indeed, it would be expected that hybrids from the same breeding programme would have a closer genetic structure, including regulatory elements, because of the admixture among genotypes sharing a common parentage. We also observed that the expression pattern of the hybrids was more similar to that of *S. officinarum* than to those of other species, suggesting preserved regulatory control of the expression of these genes after hybridization. These findings also may suggest that genotype clustering might be influenced by parental genome contributions. Indeed, all modern sugarcane varieties are hybrids between *S. officinarum* and *S. spontaneum*, followed by several backcrosses using *S. officinarum* to restore high sucrose content (Price, 1961). This breeding process resulted in the unequal contributions of each sub-genome in *Saccharum* hybrids. For instance, the genome of the R570 hybrid received approximately 80% of its chromosomes from *S. officinarum* (D’Hont, 2005), which could explain why the expression patterns of hybrids and *S. officinarum* are similar.

The observation that some OGs have tissue-specific expression may indicate that there is a sophisticated mechanism controlling their expression, perhaps mediated by TEs. Previous studies reported that TEs could potentially contribute to the emergence of new regulatory elements in a tissue-specific manner (Feschotte, 2008; Sundaram et al., 2014; Trizzino et al., 2018), which is a theory based on Barbara McClintock’s discovery that TEs can control gene expression (MCCLINTOCK, 1956). This is a plausible conjecture because more than 70% of the sugarcane genome is represented by TEs, and fragments of these elements were found in some OGs.

Evidence of OGs being regulated under stress conditions has been reported previously. Studies have demonstrated the importance of lineage-specific genes in plants subjected to biotic and abiotic stressors. For example, in cowpea (*Vigna unguiculata*), OGs seem to be more involved in drought adaptation than conserved genes, as OGs were highly induced compared to conserved genes under drought conditions (Li et al., 2019). Similarly, the expression levels of two OGs, *CpCRP1* and *CpEDR1*, are modulated when individuals of a model plant, widely studied for understanding the mechanism of desiccation, are subjected to dehydration and rehydration processes (Giarola et al., 2015). Here, most sugarcane OGs were differentially expressed in experiments in which plants were exposed to abiotic stress. However, we cannot confirm that all these genes changed their expression pattern as an adaptive response to these stresses. Further investigation needs to be carried out to experimentally validate the effectiveness of these genes for minimizing stress effects. Even though these findings were not experimentally validated, to confirm that these genes were DE, we observed a significant number of OGs that were up- and downregulated simultaneously in independent genotypes and analyses in the same experiment.

The functional prediction of OGs based on sequence homology is not possible. Additionally, most OGs have no sequence similarity to any known functional domain. Hence, we built a co-expression network to provide some clues about the functionality of these genes. This prediction method follows the guilt-by-association rationale (Zhang and Horvath, 2005), where the functional enrichment of modules containing genes co-expressed with OGs may suggest the potential biological roles of those OGs. In general, we observed OGs distributed in several network modules, indicating that these genes are involved in diverse biological activities. This is in accordance with previous studies that described OGs as being involved in several biological networks (Beike et al., 2015; Giarola et al., 2015; Khraiwesh et al., 2015; Schlötterer, 2015). Accordingly, the main biological function attributed to these genes, stress response, is itself a complex mechanism involving multiple biological pathways (Shulaev et al., 2008; Mantri et al., 2012). We highlight the evidence that most modules containing OGs were functionally enriched in biological processes related to stimulus responses, including those associated with various stresses (*Supplementary Figure 5*). In addition to canonical terms related to stresses, such as defence responses and responses to stimuli, we also detected modules enriched, for example, with genes associated with sulfate transport and responses to auxin. Genes functionally related to these processes play an important role in stress responses (Rahman, 2013; Chan et al., 2013; Shani et al., 2017).

To date, the potential biotechnological applications of a few OGs have been tested. The *QQS* gene, an Arabidopsis gene involved in carbon and nitrogen allocation, was introduced into the soybean genome and increased starch and protein levels in the leaves (Li and Wurtele, 2015; Li et al., 2015; O’Conner et al., 2018). OGs were also validated via gene editing as vital for soluble sugar metabolism in brassicas (Jiang et al., 2020).

Despite abundant evidence that these OGs are regulated across sugarcane tissues and genotypes, it is still unknown how many of these OGs are functional and produce stable proteins (Schlötterer, 2015; McLysaght and Hurst, 2016). Furthermore, although OGs are not essential for survival, they may play an important role in the response to environmental stresses (Arendsee et al., 2014; Beike et al., 2015; Li et al., 2019; Ma et al., 2020). Thus, the sugarcane OGs that we observed to be regulated under biotic and abiotic stresses may deserve special attention in future investigations to determine their biological roles.

## AUTHOR CONTRIBUTIONS

CBCS, MCM and DS: Conducted the experiments; CCBS and AA: Analysed the data; CBCS: Wrote the manuscript. All authors discussed the data, interpreted the results, read and edited the manuscript, and approved the final version.

## ACKNOWLEDGMENTS

The authors gratefully acknowledge the Fundação de Amparo à Pesquisa do Estado de São Paulo (FAPESP), the Conselho Nacional de Desenvolvimento Científico e Tecnológico (CNPq), and the Coordenação de Aperfeiçoamento de Pessoal de Nível Superior (CAPES) for financial support and fellowships.

## DATA AVAILABILITY STATEMENT

All the datasets used in this study are publicly available in the NCBI and Phytozome databases. Detailed information about the RNA-Seq experiments is listed in Supplementary Table 1.

