## Supplementary Material for "Taxonomically Restricted Genes are Associated with Responses to Biotic and Abiotic Stresses in Sugarcane (Saccharum spp.)"

### 1 Supplementary Figures

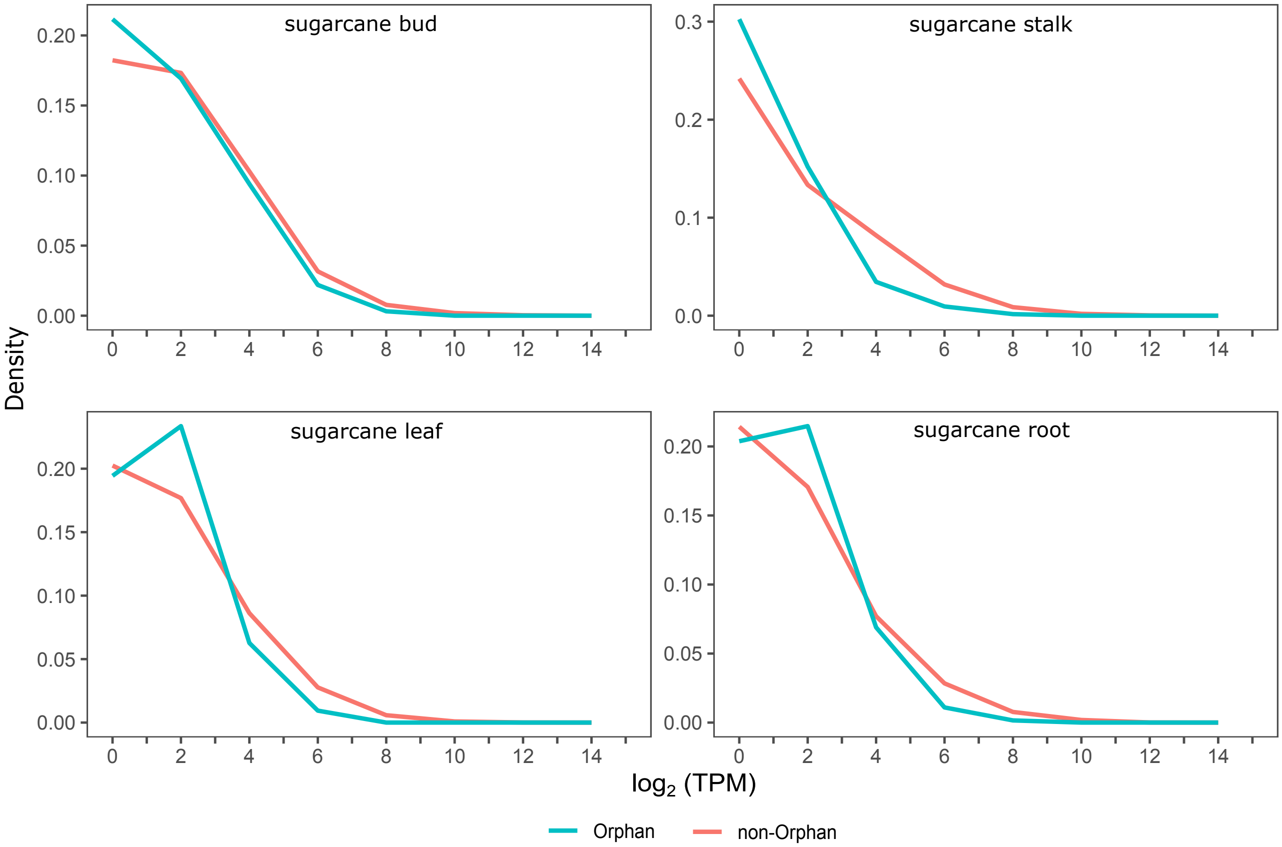


**Supplementary Figure 1.** Comparison of the expression level of orphan genes (OGs) and non-orphan genes (non-OGs) in four tissues of sugarcane.

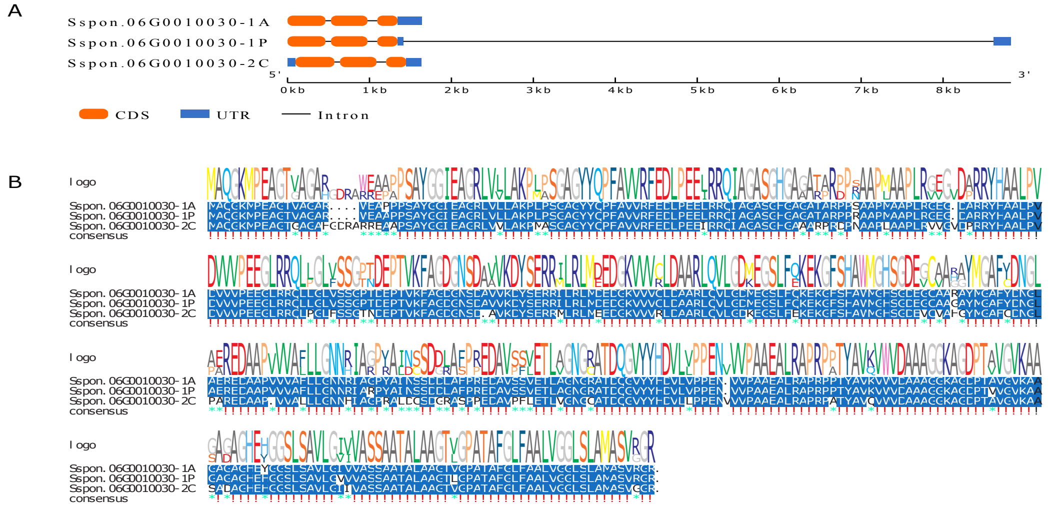


**Supplementary Figure 2**. Structure (A) and conserved motif (B) of an orphan gene (OG) induced under cold stress in both genotypes (ROC22 and Guitang08-1180). Three copies of this gene were annotated in the *S. spontaneum* genome.

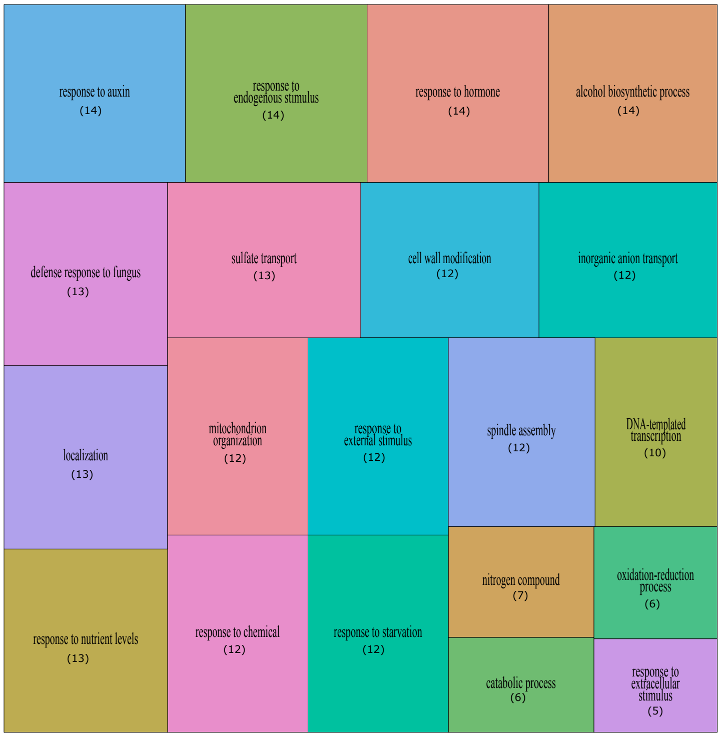


**Supplementary Figure 3**. Most frequent gene ontology (GO) terms linked to modules containing orphan genes (OGs). GO terms were selected based on enrichment analysis applying a p-value threshold of 0.05. The values inside the parentheses correspond to the number of OGs associated with these terms.
